# The Fas–FADD–caspase-8 axis is a cancer cell-intrinsic determinant of cytotoxic lymphocyte–mediated killing

**DOI:** 10.64898/2026.06.14.732110

**Authors:** Elise Solli, Shixiong Wang, Qian Wei, Nathaniel Edward Bennett Saidu, Kjetil Taskén, Youxian Li

## Abstract

Cytotoxic lymphocytes induce cancer cell death through death receptor–ligand interactions and the perforin–granzyme pathway. These pathways are generally thought to converge on the activation of executioner caspases to drive apoptosis. Here, we employed a reductionist approach to systematically disrupt key cell death mediators in a cytotoxic lymphocyte killing system to define their roles in determining cancer cell fate. We found that loss of executioner caspases conferred only limited resistance to cytotoxic lymphocyte–mediated killing. To identify cancer cell–intrinsic regulators that function beyond executioner caspases, we performed unbiased genome-wide CRISPR screens in executioner caspase-deficient cells. Unexpectedly, disruption of Fas or FADD—core components of the death receptor pathway— conferred substantial resistance to cytotoxic lymphocyte–mediated killing even in the absence of executioner caspases. This resistance persisted following additional disruption of known downstream mediators of Fas–FADD–caspase-8 (CASP8) signaling. Together, these findings identify the Fas–FADD–CASP8 axis as a central cancer cell–intrinsic determinant of susceptibility to cytotoxic lymphocyte–mediated killing whose function is not fully explained by canonical apoptotic or non-apoptotic effector pathways. Our results further suggest that CASP8 engages additional downstream substrates or mechanisms to promote cytotoxic lymphocyte–induced cancer cell death.

## Introduction

Cytotoxic lymphocytes, including cytotoxic T lymphocytes (CTLs) and natural killer (NK) cells, play fundamental roles in the elimination of cancer cells. These immune effector cells recognize malignant targets through distinct mechanisms: CTLs detect tumor–derived antigens presented on major histocompatibility complex (MHC) molecules ^1^, whereas NK cells sense aberrant expression of activating and inhibitory ligands ^2^. However, both CTLs and NK cells induce target cell death primarily through two major pathways ^3,4^. The first is the death ligand–death receptor pathway, most prominently mediated by the interaction of Fas ligand (FasL) expressed on CTLs with the Fas receptor on target cells. The second pathway involves the delivery of cytotoxic granzymes into target cells through perforin–dependent pore formation in the target cell plasma membrane. The ability of cytotoxic lymphocytes to selectively eradicate malignant cells underpins modern cancer immunotherapy, including chimeric antigen receptor (CAR) T-cell therapy, T-cell receptor (TCR)-engineered T-cell therapy, and NK cell–based therapeutic approaches ^5–7^.

A key downstream consequence of both the death receptor and perforin–granzyme pathways is the activation of apoptosis, a well-characterized form of programmed cell death mediated by caspases ^8^. Among these, caspase-3 (CASP3) serves as a principal executioner caspase in apoptosis ^9,10^. It can be activated directly by granzyme B ^11^ or via caspase-8 (CASP8) downstream of Fas–FADD signaling ^12,13^. In addition, both CASP3 and granzyme B cleave BID into truncated BID (tBID) ^14–16^, which promotes mitochondrial outer membrane permeabilization (MOMP) through BAX and/or BAK ^17,18^. This process results in the release of cytochrome c and Smac/DIABLO, facilitating apoptosome formation and caspase-9 (CASP9) activation, which ultimately leads to CASP3 activation ^19–22^. Activated CASP3 executes the apoptotic program by cleaving numerous downstream substrates ^9^, resulting in hallmark features such as apoptotic body formation and DNA fragmentation ^8,23^. Two other executioner caspases, caspase-6 (CASP6) and caspase-7 (CASP7), also exist. CASP3 and CASP7 show functional redundancy in driving apoptosis in response to both extrinsic and intrinsic apoptotic stimuli, whereas CASP6 has no apparent role in these processes ^24^.

In addition to apoptosis, alternative forms of cell death have been implicated in cytotoxic lymphocyte–mediated killing of cancer cells ^25^. Pyroptosis, a lytic form of programmed cell death mediated by pore-forming gasdermin proteins ^26,27^, can be induced in target cancer cells by cytotoxic lymphocytes ^28,29^. For example, gasdermin E (GSDME) can be cleaved and activated by CASP3 ^27,30^; therefore, in GSDME-expressing cancer cells, CASP3 can mediate both apoptosis and pyroptosis via the CASP3–GSDME axis. Gasdermins have also been reported as direct substrates of granzymes ^28,31^. Furthermore, in cases where CASP8 is inhibited, death receptor signaling can initiate necroptosis, an alternative signaling pathway mediated by receptor-interacting protein kinase 1–receptor-interacting protein kinase 3–mixed lineage kinase domain-like (RIPK1–RIPK3–MLKL) axis ^32^. Ferroptosis is another form of regulated necrotic cell death driven by plasma membrane lipid peroxidation and has also been proposed to contribute to cytotoxic lymphocyte–mediated cancer cell death ^33,34^

Despite our increased understanding of the various forms of cell death and their associated pathways, the contribution of individual pathways and mediators to cancer cell survival and resistance remains poorly understood. Indeed, although executioner caspases play critical roles in apoptosis and are essential for apoptotic hallmarks such as DNA fragmentation, they have often been reported to be dispensable for cell death itself ^15,35–37^. In such cases, cell death is frequently attributed to alternative mechanisms, including pyroptosis, necroptosis, and mitochondrial damage resulting from MOMP ^31,38^. However, a systematic evaluation of the relative contributions of alternative cell death pathways and their mediators has been lacking.

Here, we employed an unbiased approach to identify cancer cell–intrinsic factors that determine susceptibility to cytotoxic lymphocyte–mediated killing. We found that cancer cells deficient in Fas–FADD–CASP8 signaling are markedly more resistant to cytotoxic lymphocyte– mediated killing, even in the absence of executioner caspases, necroptosis, or BID–mediated mitochondrial damage. Our findings establish Fas–FADD–CASP8 signaling as a central cancer cell–intrinsic determinant of cell fate under immune attack. Furthermore, our results strongly indicate the existence of additional, as yet unidentified substrates downstream of Fas–FADD– CASP8 that contribute to cytotoxic lymphocyte–induced cancer cell death.

## Results

### Loss of executioner caspases confers minimal resistance to CTL–mediated cytotoxicity

To systematically assess the contribution of various death mediators to cytotoxic lymphocyte– mediated cancer cell death, we established a simple and robust killing system using the mouse mastocytoma cell line P815 as the target cancer cell line (**Figure 1A**). P815 cells endogenously express mouse Fcγ receptors (FcγRs) ^39^, enabling efficient recognition and killing by primary human CTLs upon coating with a mouse-derived anti-human CD3 antibody (αCD3), without the need for antigen–TCR engineering. Their rapid growth also facilitates efficient generation of knockout cell lines. In addition, P815 cells express *Casp3* but not *Casp7* (**Figure 1B**), allowing complete disruption of apoptosis through deletion of *Casp3* alone. P815 cells express *Gsdme* and are susceptible to CASP3–GSDME–mediated pyroptosis, as evidenced by GSDME cleavage (**Figure 1C**) and the characteristic ballooning morphology observed following treatment with raptinal (**Figure 1D, left panel**), a potent caspase activator. Deletion of *Casp3* (**Figure 1E**) renders the cells resistant to both apoptosis and pyroptosis induced by raptinal (**Figure 1D, right panel**). To investigate the extent to which *Casp3* deletion confers resistance to CTL–mediated killing, we expressed ZsGreen1 fluorescence in WT and *Casp3^−/−^* P815 cells and co-cultured them with primary human CTLs in the presence of αCD3. Because dead cells eventually lose fluorescence due to cessation of protein expression, quantification of the green fluorescent area provides a reliable measure of cancer cell growth (increase in green area) and death (decrease in green area) in co-culture systems. αCD3-directed killing by human CTLs efficiently eliminated WT P815 cells at an effector-to-target ratio of 1:1, as indicated by the loss of green fluorescence in co-culture within 72 hours (**Figure 1F**). Interestingly, although *Casp3^−/−^* P815 cells showed a slight delay in elimination, the majority of the cells were ultimately efficiently killed (**Figure 1F**), indicating that *Casp3* deletion confers minimal protection against CTL–mediated killing.

**Figure 1.**
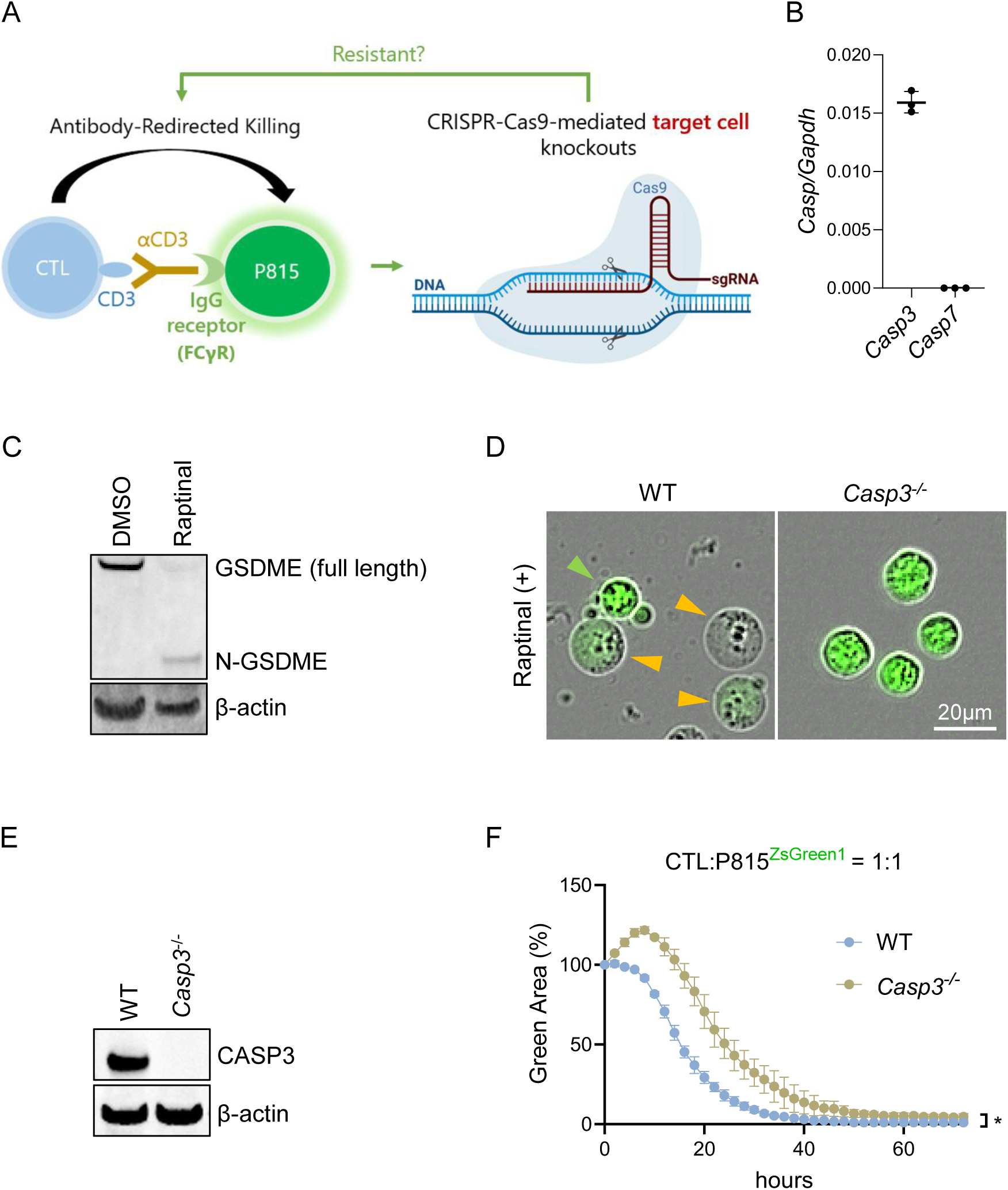
Establishment and validation of a cytotoxic T lymphocyte (CTL)–P815 killing system. **A)** Schematic overview of the CTL–P815 killing system used in this study. **B)** Transcript levels of Caspase-3 (*Casp3*) and Caspase-7 (*Casp7*) in P815 wild-type (WT) cells were quantified by reverse transcription quantitative PCR (RT-qPCR) and normalized to *Gapdh* expression. **C)** P815 WT cells were treated with the caspase activator raptinal or DMSO as a vehicle control to assess their capacity to undergo CASP3–gasdermin E (GSDME)–dependent pyroptosis. Pyroptotic induction was confirmed by GSDME cleavage into its N-terminal fragment (N-GSDME) and by characteristic ballooning morphology (see panel D, left) **D)** WT and *Casp3^−/−^*P815^ZsGreen1^ cells were treated with raptinal for 1 h to evaluate susceptibility to apoptosis and pyroptosis. Green and orange arrows indicate apoptotic and pyroptotic cells, respectively. **E)** Complete knockout (KO) of *Casp3* in P815 cells was validated by Western blot analysis. **F)** WT and *Casp3^−/−^* P815^ZsGreen1^ cells were co-cultured with activated primary human CD8⁺ CTLs (in the presence of anti-CD3) at an effector-to-target ratio of 1:1. Target cell death was quantified by loss of ZsGreen1 fluorescence and reported as the percentage of green area, normalized to the 0 h time point. *P < 0.05, two-tailed unpaired *t*-test. Data are presented as mean ± SD of three technical replicates and are representative of at least two independent experiments (F).

### Genome-wide CRISPR screening identifies Fas–FADD signaling as a key mediator of CASP3–independent CTL–mediated killing

To gain insight into cytotoxic lymphocyte–mediated, CASP3–independent cancer cell death, we leveraged the CTL–αCD3–P815 killing system and performed a genome-wide CRISPR screen. *Casp3^−/−^* P815 cells were transduced with a mouse genome-wide guide RNA (gRNA) library (Brie library; 19,674 genes targeted by four gRNAs per gene, together with 1,000 non-targeting controls; 78,637 gRNAs in total) ^40^ and subjected to CTL–mediated selection. Surviving cells were collected and sequenced to identify gRNAs enriched following selection. As shown in **Figure 2A**, gRNAs targeting FcγR genes (*Fcgr1* and *Fcgr3*), *Fcer1g*, which is required for murine FcγR assembly ^41^, and the transcription factor *Spi1*, which controls FcγR expression ^42,43^, were among the top hits. This is consistent with the requirement for αCD3– FcγR coupling to enable CTL–P815 engagement in this system (**Figure 1A**). Notably, despite CASP3 being a downstream effector of Fas–FADD signaling and already deleted prior to the screen, both *Fas* and *Fadd* emerged as top hits (**Figure 2A, Supplementary Table I**). In addition, *Sec62*—an endoplasmic reticulum translocation factor involved in the biogenesis of multiple secretory and transmembrane proteins ^44–46^—was also identified among the top hits (**Figure 2A, Supplementary Table I**). Flow cytometry confirmed that *Casp3^−/−^Sec62^−/−^* cells (**Figure 2B, left panel**) lacked surface expression of Fas (**Figure 2C**), further supporting the importance of Fas–FADD signaling in driving cancer cell death even in the absence of CASP3. Deletion of either *Fas* or *Fadd* (**Figure 2B, middle and right panels**) rendered *Casp3^−/−^* cells considerably more resistant to CTL–mediated killing (**Figure 2D**). Consistently, *Casp3^−/−^Sec62^−/−^* cells exhibited resistance to CTL–mediated cell death comparable to that of *Casp3^−/−^Fas^−/−^* and *Casp3^−/−^Fadd^−/−^*cells (**Figure 2E**). These data suggest that death signaling through the Fas– FADD pathway remains a major driver of CTL–mediated cancer cell death even in the absence of CASP3.

**Figure 2.**
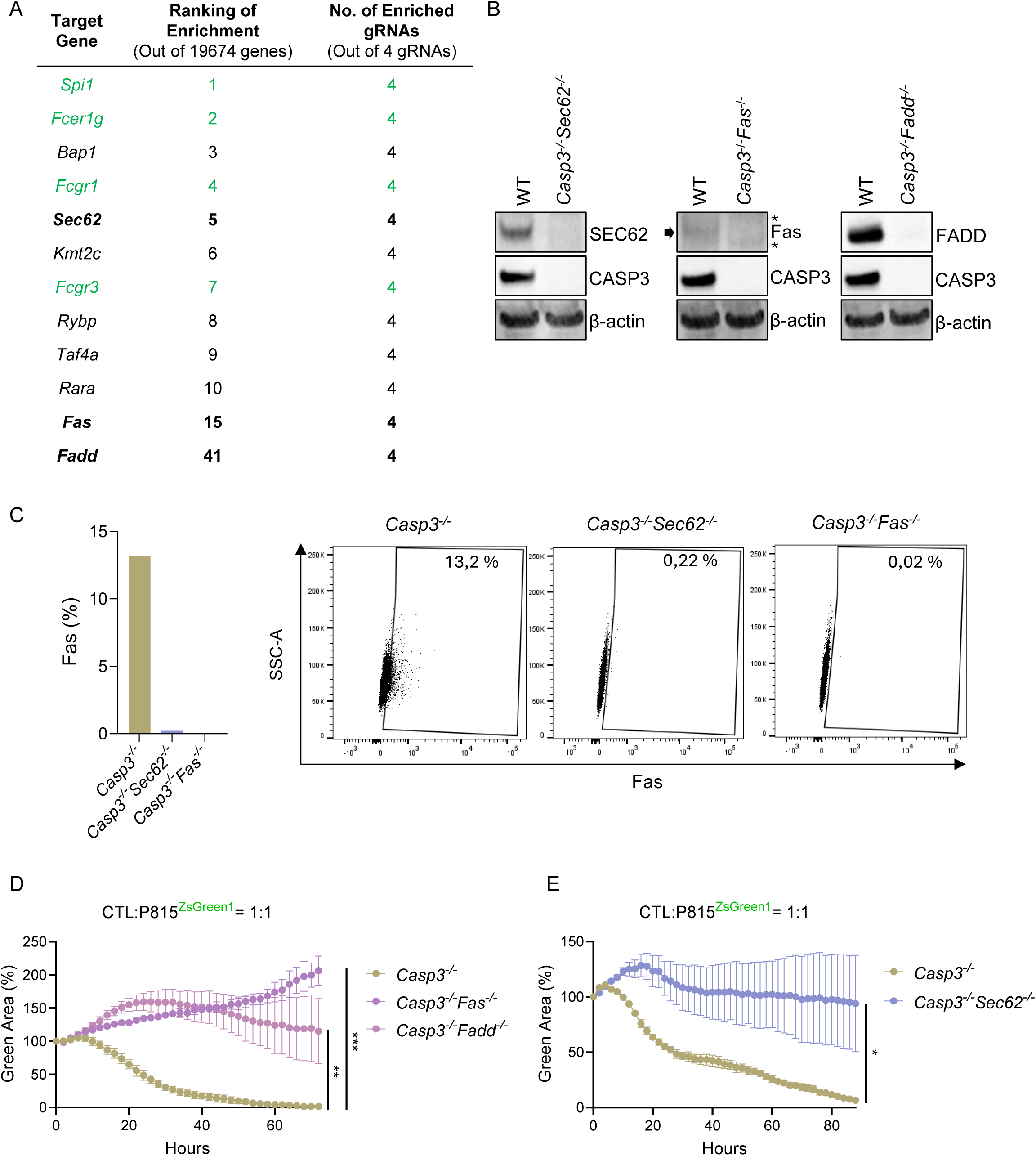
Identification and validation of candidate mediators of Caspase-3 (CASP3)– independent cell death induced by cytotoxic T lymphocytes (CTLs). **A)** Ranked list of top candidate genes identified from the genome-wide CRISPR–Cas9 screen as putative mediators of CASP3–independent cell death. Positive control genes associated with Fcγ receptor (FcγR) expression are highlighted in green. **B)** Complete knockout (KO) of *Sec62*, *Fas*, and *Fadd* in P815 *Casp3^−/−^*cells was confirmed by Western blot analysis. The arrow indicates the Fas-specific band, and the asterisks indicate two faint non-specific bands. **C)** Surface expression of Fas was assessed by flow cytometry in *Casp3^−/−^*, *Casp3^−/−^Sec62^−/−^* and *Casp3^−/−^Fas^−/−^*P815 cells. The percentage of Fas-positive cells is shown in the left panel, with flow cytometry plots shown in the right panel. D) *Casp3^−/−^*, *Casp3^−/−^Fas^−/−^*, or *Casp3^−/−^Fadd^−/−^* P815^ZsGreen1^ cells were co-cultured with activated primary human CTLs (in the presence of anti-CD3) at an effector-to-target ratio of 1:1. Target cell death was quantified by loss of ZsGreen1 fluorescence and reported as the percentage of green area, normalized to the 0 h time point. **E)** CTL–mediated killing of *Casp3^−/−^* or *Casp3^−/−^Sec62^−/−^* P815^ZsGreen1^ cells was assessed as in (D). *P < 0.05, **P < 0.01, ***P < 0.001, one-way ANOVA (D) and two-tailed unpaired *t*-test (E). Data are presented as mean ± SD of three technical replicates and are representative of at least two independent experiments (D and E).

### Necroptotic and mitochondrial pathways partially contribute to CASP3–independent CTL-mediated killing

Death receptor–ligand engagement can trigger necroptosis, an alternative form of cell death that occurs under conditions of CASP8 inhibition and is mediated by activation of the RIPK1– RIPK3–MLKL axis ^32^. Deletion of the effector caspase *Casp3* in our system may induce a negative feedback effect on upstream CASP8 activation ^24^, thereby promoting diversion of cells toward activation of the necroptotic program. To assess the contribution of necroptosis to cytotoxic lymphocyte–mediated, CASP3–independent cancer cell death, we generated a *Casp3^−/−^Mlkl^−/−^* double-knockout P815 cell line (**Figure 3A, left panel**). Deletion of *Mlkl* had a modest protective effect against CTL–mediated killing of *Casp3^−/−^*P815 cells (**Figure 3B**), suggesting that MLKL–dependent necroptosis only partially contributes to CASP3–independent cancer cell death induced by CTLs.

**Figure 3.**
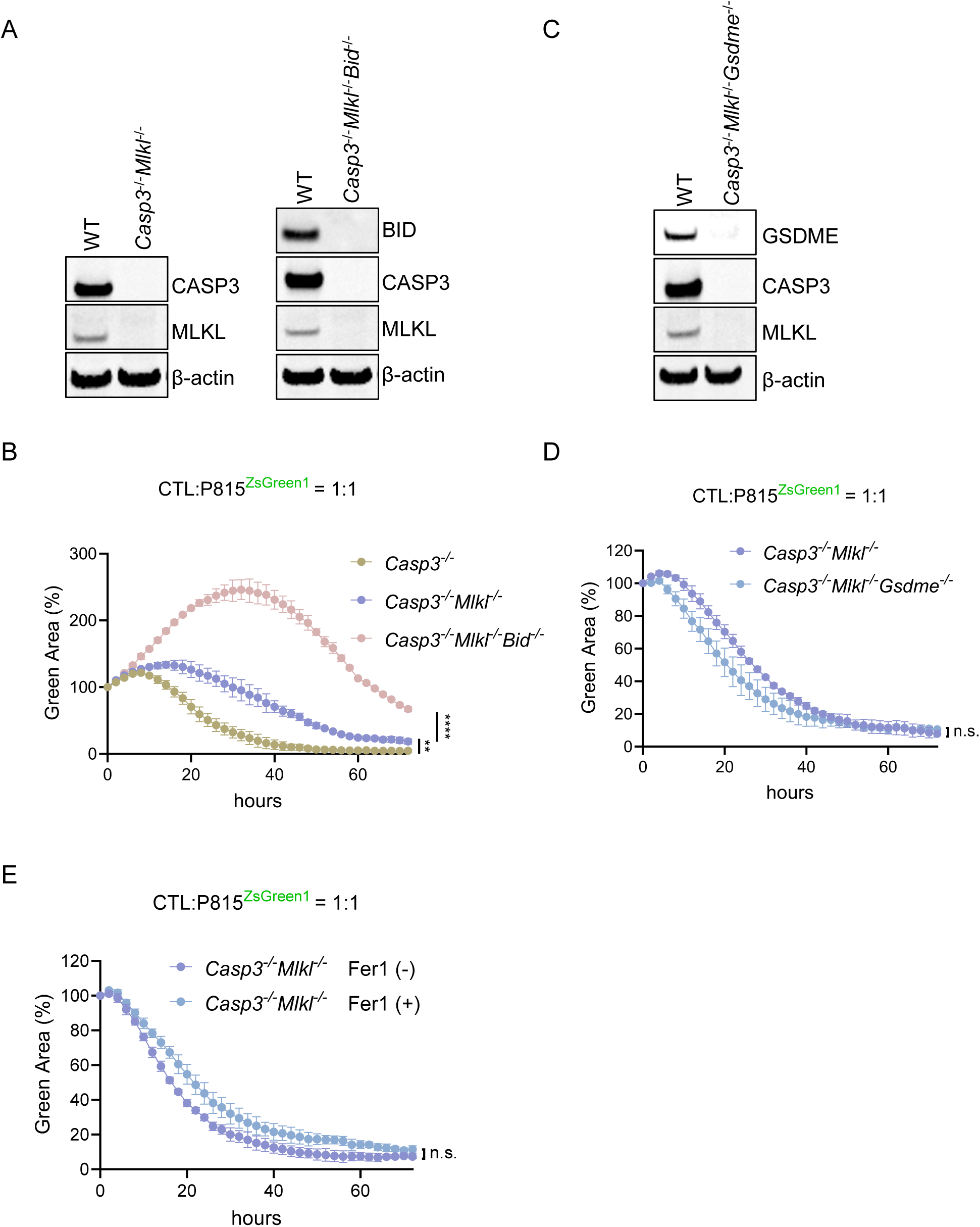
Genetic and pharmacological dissection of Caspase-3 (CASP3)–independent cytotoxic T lymphocyte (CTL)–mediated killing pathways. **A)** Western blot analysis confirming complete knockout (KO) of *Casp3* and *Mlkl* (left panel), and of *Casp3*, *Mlkl*, and *Bid* (right panel) in P815 cells. B) *Casp3^−/−^*, *Casp3^−/−^Mlkl^−/−^*, or *Casp3^−/−^Mlkl^−/−^Bid^−/−^* P815^ZsGreen1^ cells were co-cultured with activated primary human CD8⁺ CTLs (in the presence of anti-CD3) at an effector-to-target ratio of 1:1. Target cell death was quantified by loss of ZsGreen1 fluorescence and reported as the percentage of green area, normalized to the 0 h time point. **C)** Western blot analysis confirming complete KO of *Casp3*, *Mlkl*, and *Gsdme* in P815 cells. **D)** CTL–mediated killing was assessed as described in (B) using *Casp3^−/−^Mlkl^−/−^* and *Casp3^−/−^Mlkl^−/−^Gsdme^−/−^*P815 cells to evaluate the contribution of GSDME to target cell death. **E)** CTL– mediated killing was assessed as described in (B) using *Casp3^−/−^Mlkl^−/−^*P815 cells in the presence or absence of the ferroptosis inhibitor ferrostatin-1 (Fer-1) to assess the potential contribution of ferroptosis to target cell death. n.s., not significant, **P < 0.01, ****P < 0.0001, one-way ANOVA (B) and two-tailed unpaired *t*-test (D and E). Data are presented as mean ± SD of three technical replicates and are representative of at least two independent experiments (B, D, and E).

GSDME has been proposed to serve as a direct substrate of granzyme B ^31^. To assess the contribution of the putative granzyme B–GSDME axis to CTL–induced cell death, we generated a *Casp3^−/−^Mlkl^−/−^Gsdme^−/−^* P815 cell line (**Figure 3C**) and compared its susceptibility to CTL–mediated killing with that of its parental *Casp3^−/−^Mlkl^−/−^* line (**Figure 3D**). No difference in sensitivity was observed between the two cell lines, suggesting that GSDME contributes minimally to CTL–induced cell death in the absence of CASP3, at least in this experimental system. Similarly, pretreatment of *Casp3^−/−^Mlkl^−/−^* cells with ferrostatin-1, a specific inhibitor of ferroptosis, had little protective effect (**Figure 3E**), indicating that ferroptosis plays a minimal role in this context.

MOMP is often considered a point of no return because, in addition to releasing intermembrane space proteins such as cytochrome c and Smac/DIABLO into the cytosol to promote apoptosome formation and CASP9–dependent activation of CASP3, it can also induce mitochondrial injury, reactive oxygen species (ROS) production, and oxidative stress ^47,48^. To assess the contribution of BID–mediated MOMP to CASP3–independent CTL–mediated cancer cell death, we generated a *Casp3^−/−^Mlkl^−/−^Bid^−/−^*triple-knockout P815 cell line. Deletion of *Bid* in the *Casp3^−/−^Mlkl^−/−^*background (**Figure 3A, right panel**) further increased resistance to CTL–mediated killing (**Figure 3B**), suggesting that BID–mediated MOMP and mitochondrial damage also contributes to CTL–induced cell death even when downstream apoptotic execution is impaired.

### Fas–FADD–CASP8 signaling remains essential after disruption of apoptotic, necroptotic, and mitochondrial death pathways

We observed that the triple-knockout P815 cell line *Casp3^−/−^Mlkl^−/−^Bid^−/−^*initially expanded following CTL attack (**Figure 3B**) but eventually succumbed to cell death, as indicated by a transient increase followed by a reduction of ZsGreen1 signal. This observation led us to hypothesize that CTL–mediated cancer cell death cannot be fully explained by apoptosis, necroptosis, and BID–mediated mitochondrial damage alone. To gain insight into additional factors contributing to CTL–induced cancer cell death, we performed another genome-wide CRISPR screen using the newly generated *Casp3^−/−^Mlkl^−/−^Bid^−/−^* cell line. Interestingly, following CTL selection, components of the Fas–FADD signaling pathway (*Fas, Fadd,* and *Casp8*), as well as the Fas membrane expression regulator *Sec62*, again emerged as top hits (**Figure 4A, Supplementary Table II**). To validate these findings, we generated quadruple-knockout P815 cell lines (*Casp3^−/−^Mlkl^−/−^Bid^−/−^Fas^−/−^, Casp3^−/−^Mlkl^−/−^Bid^−/−^Fadd^−/−^, and Casp3^−/−^Mlkl^−/−^Bid^−/−^Casp8^−/−^*) (**Figure 4B**) and compared them with their parental *Casp3^−/−^Mlkl^−/−^Bid^−/−^*counterpart. The quadruple-knockout cells displayed markedly increased resistance to CTL–mediated killing (**Figure 4C, D**). Together, these data strongly indicate that the Fas–FADD–CASP signaling axis is a critical determinant of cancer cell fate, governing survival versus death even when apoptosis, necroptosis, and BID–mediated mitochondrial damage are impaired.

**Figure 4.**
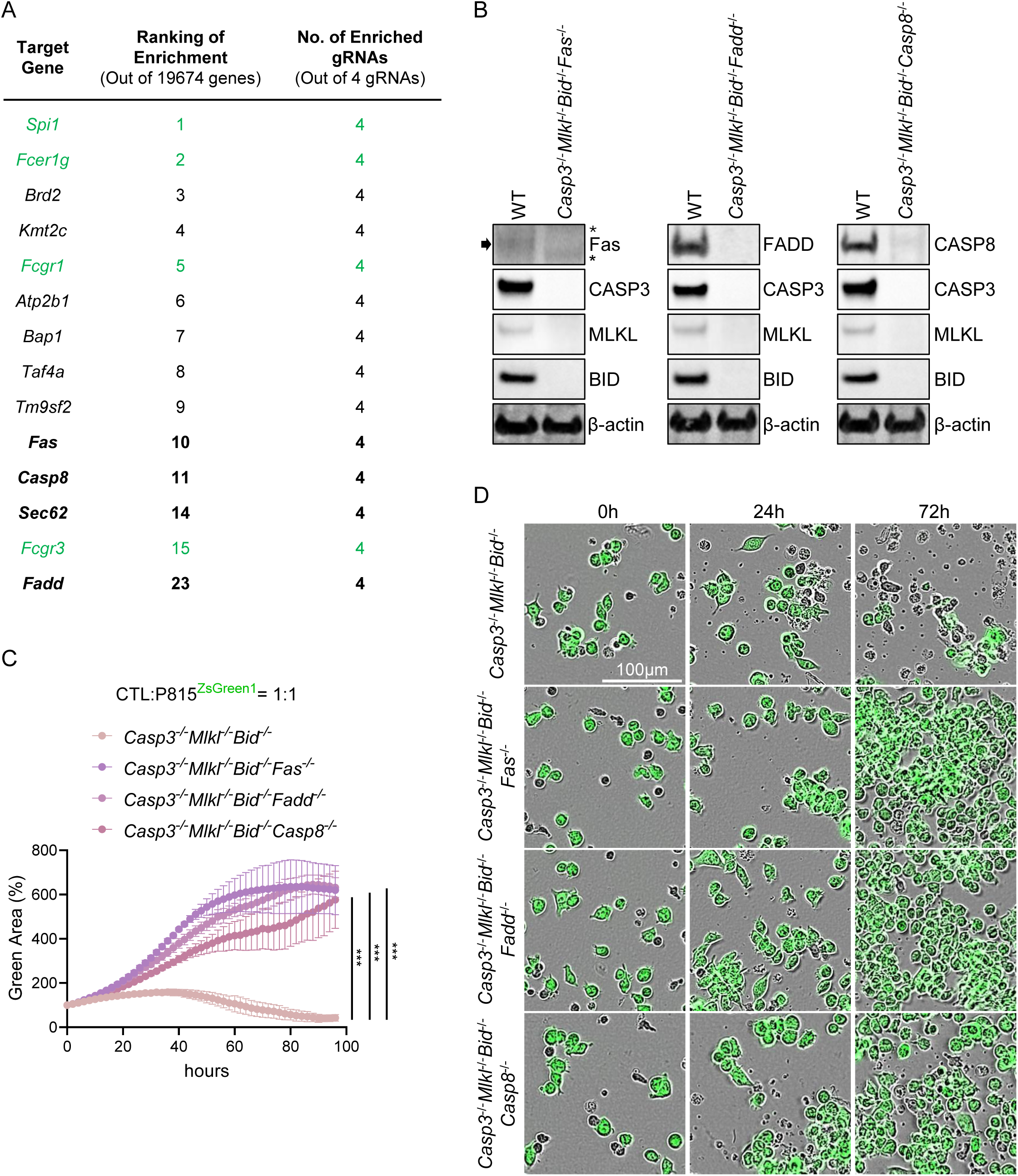
Identification of Fas–FADD–caspase-8 (CASP8) axis as a mediator of residual cytotoxic T lymphocyte (CTL)–induced killing in a CASP3/MLKL/BID–deficient background. **A)** Ranked list of top candidate genes identified from the genome-wide CRISPR– Cas9 screen performed in the *Casp3^−/−^Mlkl^−/−^Bid^−/−^* background, representing putative mediators of residual CTL-mediated killing. Positive control genes associated with Fcγ receptor (FcγR) expression are highlighted in green. **B)** Western blot analysis confirming complete knockout (KO) of *Fas*, *Fadd*, and *Casp8* in the *Casp3^−/−^Mlkl^−/−^Bid^−/−^*background. The arrow indicates the Fas-specific band, and the asterisks indicate two faint non-specific bands. C) *Casp3^−/−^Mlkl^−/−^Bid^−/−^*, *Casp3^−/−^Mlkl^−/−^Bid^−/−^Fas^−/−^*, *Casp3^−/−^Mlkl^−/−^Bid^−/−^Fadd^−/−^*, or *Casp3^−/−^Mlkl^−/−^Bid^−/−^Casp8^−/−^*P815^ZsGreen1^ cells were co-cultured with activated primary human CD8⁺ CTLs (in the presence of anti-CD3) at an effector-to-target ratio of 1:1. Target cell death was quantified by loss of ZsGreen1 fluorescence and reported as the percentage of green area, normalized to the 0 h time point. **D)** Representative Incucyte^®^ images showing the death or proliferation of the indicated P815^ZsGreen1^ cell strains (green) at the indicated time points following CTL-mediated killing (effector-to-target ratio, 1:1). ***P < 0.001, one-way ANOVA. Data are presented as mean ± SD of three technical replicates and are representative of at least two independent experiments (C).

### Fas–FADD–CASP8 signaling governs cancer cell susceptibility to NK cell–mediated cytotoxicity in HeLa cells

To validate the role of Fas–FADD–CASP8 signaling in an independent cytotoxic lymphocyte– mediated killing system, we employed a complementary reductionist model based on NK-92– mediated killing of HeLa cells, using NK-92 as an NK cell line. We first generated a *CASP3^−/−^ CASP7^−/−^* double-knockout Hela^EGFP^ cell line (**Figure 5A**) and confirmed that these cells were resistant to raptinal–induced apoptosis and pyroptosis (**Figure 5B**). We then assessed killing of wild-type and *CASP3^−/−^CASP7^−/−^* Hela^EGFP^ cells by NK-92 cells. At an effector-to-target ratio of 1:1, killing was inefficient for both cell lines (**Figure 5C, upper panel**). At a 3:1 effector-to-target ratio, deletion of CASP3 and CASP7 conferred little non-significant resistance to NK-92–mediated killing compared with wild-type cells (**Figure 5C, lower panel**). This is consistent with our observation in P815 cells that disruption of executioner caspases provides little protection against cytotoxic lymphocyte–mediated killing (**Figure 1F**). HeLa cells are known to be resistant to necroptosis due to defective RIPK3 expression ^32^; therefore, disruption of necroptosis was not performed.

**Figure 5.**
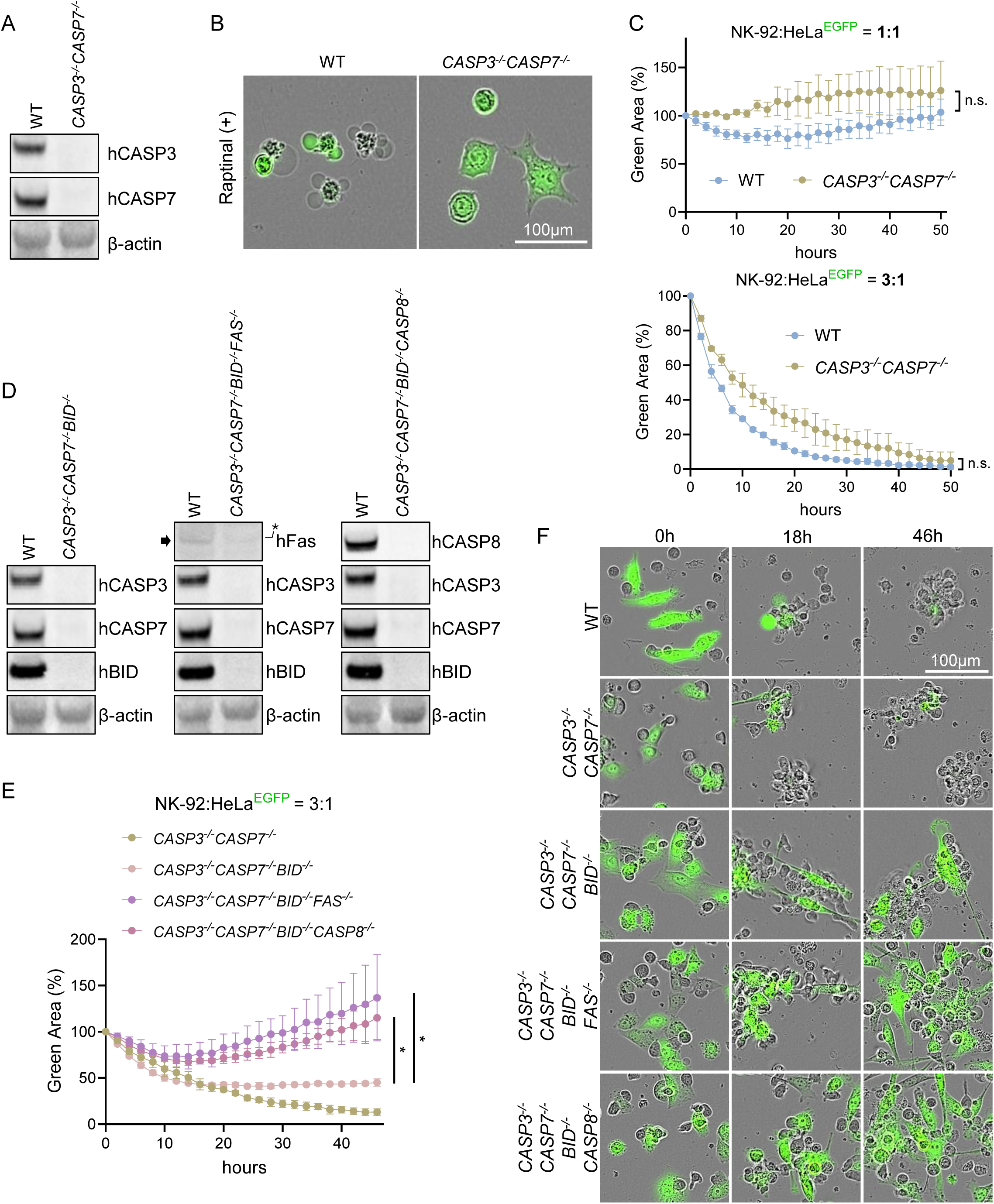
The Fas–FADD–caspase-8 (CASP8) axis governs cytotoxic lymphocyte– mediated killing in human cancer. **A)** Western blot analysis confirming complete knockout (KO) of *CASP3* and *CASP7* in HeLa^EGFP^ cells. **B)** Wild-type (WT) or *CASP3^−/−^CASP7^−/−^* HeLa^EGFP^ cells were treated with the caspase activator raptinal for 6 h to assess their capacity to undergo apoptosis and pyroptosis. Resistance to caspase-dependent apoptosis and pyroptosis in *CASP3^−/−^CASP7^−/−^* cells was confirmed by the absence of apoptotic bodies or ballooning morphology. **C)** WT or *CASP3^−/−^CASP7^−/−^* HeLa^EGFP^ cells were co-cultured with NK-92 cells, and cytotoxic killing assays were performed at the indicated effector-to-target ratio. Target cell death was quantified by loss of EGFP fluorescence and reported as the percentage of green area, normalized to the 0 h time point. **D)** Western blot analysis confirming complete knockout (KO) of *BID* in the *CASP3^−/−^CASP7^−/−^* background and of *FAS* and caspase-8 (*CASP8*) in the *CASP3^−/−^CASP7^−/−^BID^−/−^*background. The arrow indicates the Fas-specific band, and the asterisk indicates a faint non-specific band. **E)** Cytotoxic killing assays were performed as described in (C) using *CASP3^−/−^CASP7^−/−^*, *CASP3^−/−^CASP7^−/−^BID^−/−^*, *CASP3^−/−^CASP7^−/−^BID^−/−^FAS*⁻/⁻, or *CASP3^−/−^CASP7^−/−^BID^−/−^CASP8^−/−^*HeLa^EGFP^ cells. **F)** Representative Incucyte^®^ images showing the death or proliferation of the indicated HeLa^EGFP^ cell strains (green) at the indicated time points following NK-92-mediated killing (effector-to-target ratio, 3:1). n.s., not significant, *P < 0.05, two-tailed unpaired *t*-test (C) and one-way ANOVA (E). Data are presented as mean ± SD of three technical replicates and are representative of at least two independent experiments (C and E).

Consistent with our observations in P815 cells, additional deletion of *BID* (**Figure 5D, left panel**) in the *CASP3^−/−^CASP7^−/−^* background conferred resistance to NK cell–mediated killing (**Figure 5E, F**). Importantly, subsequent deletion of either *FAS* or *CASP8* (*CASP3^−/−^CASP7^−/−^ BID^−/−^FAS^−/−^*and *CASP3^−/−^CASP7^−/−^BID^−/−^CASP8^−/−^*) (**Figure 5D, middle and right panels**) rendered the cells markedly more resistant to NK-92–mediated killing than their parental *CASP3^−/−^CASP7^−/−^BID^−/−^*counterpart (**Figure 5E, F**). Together, these findings further establish the Fas–FADD–CASP8 signaling pathway as a critical cancer cell–intrinsic determinant of cytotoxic lymphocyte–induced cell death, operating in both CTL– and NK cell–mediated killing.

### CASP8 is preferentially mutated in cancer compared with known downstream substrates

The importance of the FAS–FADD–CASP8 signaling pathway in conferring cancer resistance is underscored by the substantially higher frequency of *CASP8* mutations observed in cancer patients compared with mutations in its downstream effectors. Analysis of The Cancer Genome Atlas (TCGA) data identified simple somatic mutations (SSMs) in *CASP8* in 1.91% of patients across all cohorts (349 of 18,289 cases) (**Figure 6A**). In contrast, SSMs in *CASP3*, *CASP7*, and *BID* occurred at considerably lower frequencies, with 74, 93, and 77 cases, respectively, among the 18,289 patients analyzed (**Figure 6A**). Moreover, *CASP8* also harbored a markedly greater number of distinct SSMs, with 286 unique mutations compared with 66, 80, and 74 for *CASP3*, *CASP7*, and *BID*, respectively (**Figure 6B**). The necroptosis effector *MLKL* exhibited slightly higher SSM frequencies and a greater number of distinct SSMs than *CASP3*, *CASP7*, and *BID* (**Figure 6A, B**). Interestingly, *FAS* and *FADD* displayed lower SSM frequencies than *CASP8*, suggesting that *CASP8* is more susceptible to genetic alteration within this signaling axis (**Figure 6A, B**). The Ensembl Variant Effect Predictor (VEP) ^49^ indicated that a substantial proportion (26%) of *CASP8* SSMs are predicted to have high-impact (disruptive) consequences, including protein truncation, loss of function, or nonsense-mediated decay. Notably, *CASP8* has the highest proportion of high-impact variants among the seven genes examined (**Figure 6C, D).** Further analysis of the most recurrent SSMs (present in ≥5 of the 18,289 cases) in *CASP8* revealed that the majority are likely loss-of-function mutations, with six of the nine most frequent variants predicted to cause frameshift or premature stop codons (**Figure 6E**). These clinical observations are consistent with our experimental findings that the Fas–FADD–CASP8 signaling axis plays a central role in determining cancer cell susceptibility to cytotoxic lymphocyte–mediated killing. They further support the notion that this pathway has functions beyond its established roles in canonical cell death and suggest that disruption of *CASP8* provides a selective advantage during cancer evolution.

**Figure 6.**
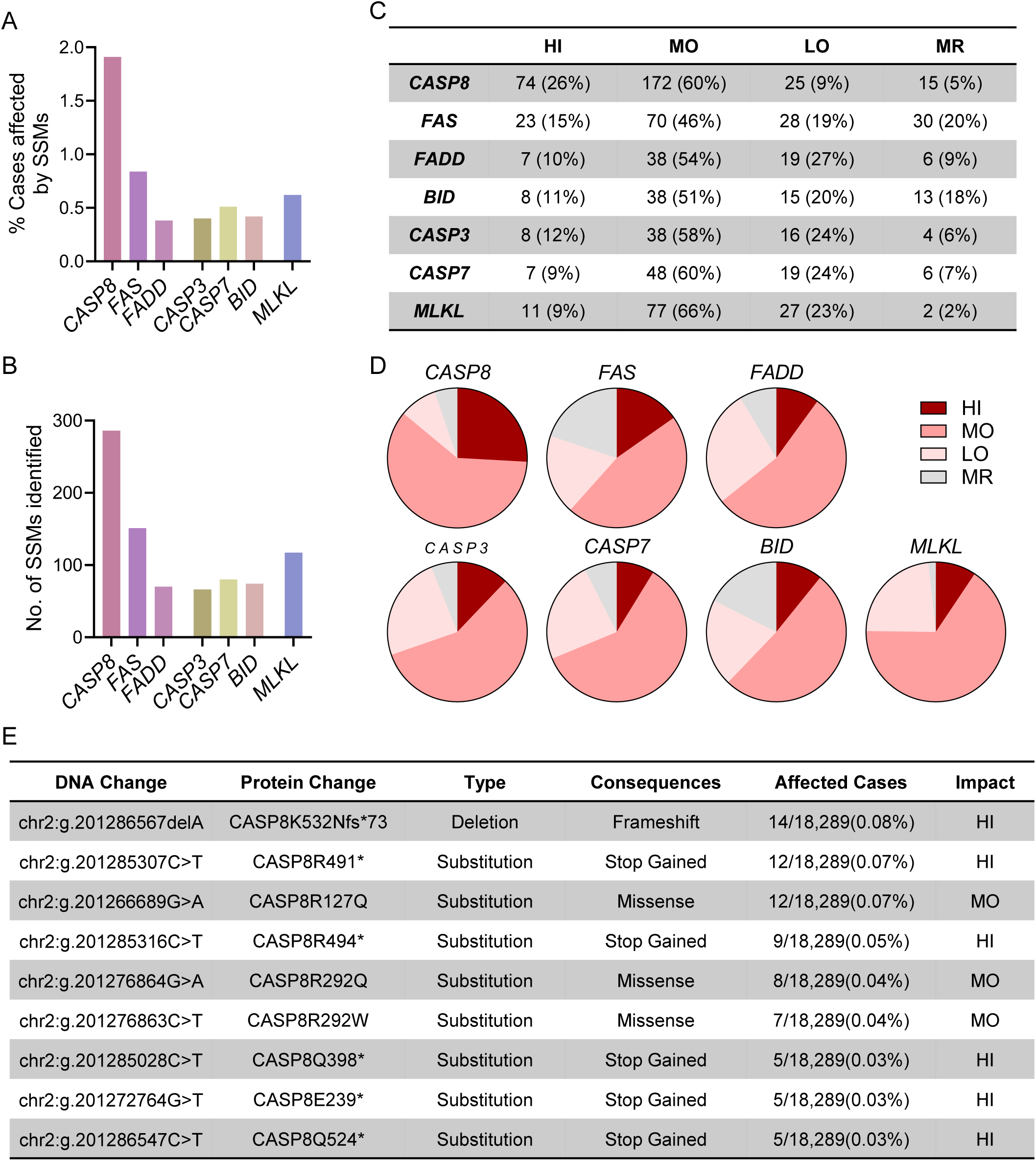
CASP8 is preferentially mutated in human cancers. **A–B)** The number of cancer cases across all cancer types (pan-cancer; total n = 18,289) harboring simple somatic mutations (SSMs) in selected cell death regulators (A), and the total number of distinct SSMs identified per gene (B), were retrieved from The Cancer Genome Atlas (TCGA). **C–D)** Distinct SSMs for each gene were categorized according to predicted functional impact based on Ensembl Variant Effect Predictor (VEP) annotations (HI, high impact; MO, moderate impact; LO, low impact; MR, modifier). Definitions of impact categories are provided in the Materials and Methods. **E)** Most frequent SSMs in *CASP8*, defined as variants detected in ≥5 of the 18,289 cases.

## Discussion

In this study, we used targeted genetic disruption to evaluate the contribution of multiple known death mediators involved in apoptosis, pyroptosis, necroptosis, and MOMP to cytotoxic lymphocyte–induced cancer cell death. In parallel, we performed unbiased genome-wide CRISPR screens to systematically identify cancer cell–intrinsic factors that determine survival or death following cytotoxic lymphocyte attack. Compared with the most reductionist experimental approaches using recombinant death ligands or granzymes, our co-culture killing system more closely recapitulates the biological context of cytotoxic lymphocyte–mediated killing of cancer cells, in which multiple effector mechanisms act in parallel on target cells. In addition, we focused on perturbation of cancer cell–intrinsic factors rather than effector cell components (e.g., perforin or granzyme expression in cytotoxic lymphocytes), based on the rationale that cancer cells may exploit their intrinsic genetic programs to evade immune attack. Our screens consistently identified the death receptor–ligand pathway, signaling through the Fas–FADD–CASP8 axis, as a key determinant of cancer cell fate. We validated the importance of this pathway in two independent killing systems using CTLs or NK cells as effector populations.

Various methodologies have been developed to measure cell death, and care must be taken when interpreting their readouts, as many approaches have intrinsic limitations. For example, CASP3/7 substrates are frequently used as indicators of cancer cell death; however, this approach inherently detects only apoptosis or CASP3–GSDME–mediated pyroptosis and therefore excludes other forms of cell death. Similarly, methods that rely on membrane permeabilization, such as calcein release assays, lactate dehydrogenase (LDH) release assays, and intracellular DNA staining, tend to emphasize necrotic forms of death, because plasma membrane integrity is generally preserved during apoptosis, at least in its early stages ^50^. Viability assays based on ATP quantification or cellular metabolic activity, such as CellTiter-Glo and MTT assays, are difficult to apply in co-culture systems because the presence of effector immune cells confounds interpretation of the measurements. In this study, we adopted an unbiased approach in which cancer cell death or expansion was quantified based on endogenously expressed fluorescent signals. Because dead cancer cells cease fluorescent protein expression and eventually lose fluorescence, whereas surviving cells continue to proliferate, this strategy provides a more objective assessment of overall cancer cell survival and death.

Although the extrinsic death signaling pathway mediated by Fas–FADD–CASP8 has been recognized for many years, its importance has primarily been appreciated in the context of apoptosis through downstream activation of the executioner caspases CASP3 and CASP7 ^51,52^, either directly by CASP8 or indirectly via CASP8–mediated cleavage of BID ^53^. Our data suggest that the Fas–FADD–CASP8 axis is more broadly important than previously appreciated, remaining functionally relevant even when canonical downstream substrates, including CASP3, CASP7, and BID, are fully disrupted.

Notably, CASP3, despite being a central mediator of apoptosis and an upstream regulator of GSDME–mediated pyroptosis, is rarely mutated in cancer ^54,55^. This somewhat counterintuitive observation raises the possibility that executioner caspases are not the ultimate determinants of cell death per se but rather function as downstream effectors that channel cells into specific forms of death following activation of upstream death signals. Caspase–independent cell death has long been recognized ^35^, and MOMP has been proposed to represent a point of no return for cell death, irrespective of downstream caspase activation ^48^. Nevertheless, CTLs have been shown to abrogate target cell proliferation and induce cell death even upon inhibition of both executioner caspases and mitochondrial events ^37^. This finding is consistent with our observation that systematic disruption of several well-established mediators of cell death— including *CASP3*/*7* (apoptosis and pyroptosis), *MLKL* (necroptosis), and *BID* (MOMP)—did not fully prevent cytotoxic lymphocyte–induced killing. Importantly, even when these downstream death mediators were disrupted, additional deletion of upstream Fas–FADD– CASP8 components conferred substantial resistance to cytotoxic lymphocyte–mediated killing. Consistent with this, analysis of TCGA data shows that alterations in *CASP8* are more frequently observed in cancer than mutations in *CASP3*, *CASP7*, or *BID*. It is noteworthy that CASP3, CASP7, and BID can also be activated by granzyme B, meaning that loss of functional CASP8 does not completely abolish apoptosis or CASP3–GSDME-mediated pyroptosis induced by cytotoxic lymphocytes. Furthermore, CASP8 deficiency has the potential to promote necroptosis in cancer cells. Therefore, the relatively high frequency of CASP8 mutations observed in human cancers suggests that CASP8 inactivation provides selective advantages that cannot be fully explained by currently recognized cell death mechanisms alone. Together, our findings from cytotoxic lymphocyte killing assays, genome-wide CRISPR screens, and clinical observations strongly indicate that additional, as yet unidentified, CASP8 substrates or downstream effectors contribute to cytotoxic lymphocyte–induced cancer cell death.

In summary, using a reductionist genetic approach, we provide compelling evidence that the Fas–FADD–CASP8 signaling pathway governs cancer cell susceptibility to cytotoxic lymphocyte–mediated killing. The relevance of this pathway extends beyond its established roles in apoptosis, necroptosis, and mitochondrial damage. Our work offers a potential explanation for why certain well-established death mediators, such as *CASP3*, are rarely mutated in cancer. These findings also highlight the importance of future studies aimed at identifying previously unrecognized CASP8 substrates that may contribute to cancer cell death beyond currently defined canonical cell death pathways.

## Materials and Methods

### Cell lines and cell culture

The mouse mastocytoma cell line P815 and the human natural killer cell line NK-92 were obtained from ATCC. The Lenti-X™ 293T cell line was obtained from Takara Bio. P815 cells used in this study were transduced with pLVX-IRES-ZsGreen1 (V012720, NovoPro) to constitutively express ZsGreen1 fluorescence. HeLa^EGFP^ cells were obtained from Cell Biolabs. P815, HeLa^EGFP^, and Lenti-X™ 293T cells were cultured in Dulbecco’s Modified Eagle Medium (DMEM) supplemented with 10% fetal bovine serum (FBS; Gibco) and 1% penicillin-streptomycin (Sigma-Aldrich, St. Louis, MO, USA). NK-92 cells were cultured in MEM α (Gibco) supplemented with 12.5% horse serum (Gibco), 12.5% FBS (Gibco), 2 mM L-glutamine (Gibco), 1% penicillin-streptomycin (Sigma-Aldrich), and 200 U/mL recombinant human IL-2 (R&D Systems). All cell cultures were maintained at 37 °C in a humidified atmosphere containing 5% CO₂.

### Isolation and activation of human T cells

Human T cells were isolated from buffy coats obtained from healthy anonymous blood donors through the Oslo University Hospital Blood Centre. Sample collection and use were approved by the Regional Committee for Medical and Health Research Ethics (REK approval no. 280751). Total T cells were isolated using the RosetteSep™ Human T Cell Enrichment Cocktail (STEMCELL Technologies), followed by density gradient centrifugation using Lymphoprep™ (STEMCELL Technologies), according to the manufacturer’s instructions.

CD8⁺ CTLs were subsequently isolated from total T cells using Dynabeads™ CD8 (Invitrogen) according to the manufacturer’s instructions. Purified CD8⁺ T cells were activated using Dynabeads™ Human T-Activator CD3/CD28 (Gibco) and cultured for 7–15 days in RPMI 1640 medium containing L-glutamine (VWR), supplemented with 10% FBS (Gibco), 1% non-essential amino acids (Gibco), 1% sodium pyruvate (Gibco), 1% penicillin-streptomycin (Sigma-Aldrich), and 100 U/mL recombinant human IL-2 (R&D Systems). Cells were maintained at 37 °C in a humidified atmosphere containing 5% CO₂.

### Reverse transcription quantitative PCR (RT-qPCR)

A total of 3 × 10⁵ cells were harvested, and total RNA was isolated using the RNeasy® Mini Kit (Qiagen,) according to the manufacturer’s instructions. Complementary DNA (cDNA) was synthesized from the isolated RNA using the RevertAid First Strand cDNA Synthesis Kit (Thermo Fisher Scientific) following the manufacturer’s protocol.

RT-qPCR was performed using PowerTrack™ SYBR™ Green Master Mix (Thermo Fisher Scientific) according to the manufacturer’s instructions. Forward and reverse primers for the target genes were purchased from Sigma-Aldrich. Primer sequences used in this study were as follows:

*Casp3*

- Forward: 5′-GGAGTCTGACTGGAAAGCCGAA-3′
- Reverse: 5′-CTTCTGGCAAGCCATCTCCTCA-3′

*Casp7*

- Forward: 5′-CCGTCCACAATGACTGCTCTTG-3′
- Reverse: 5′-CCCGTAAATCAGGTCCTCTTCC-3′

*Gapdh*

- Forward: 5′-CATCACTGCCACCCAGAAGACTG-3′
- Reverse: 5′-ATGCCAGTGAGCTTCCCGTTCAG-3′

### Western Blot

For Western blot analysis, cells were harvested and lysed in Triton™ X-100 Lysis Buffer (Thermo Fisher Scientific) supplemented with Halt™ Protease Inhibitor Cocktail (Thermo Fisher Scientific). Cell lysates were prepared at a concentration of 1 × 10⁶ cells/100µL and centrifuged at 15,000 × *g* for 10 min at 4°C. The supernatants were collected and mixed with Invitrogen™ NuPAGE™ LDS Sample Buffer and Invitrogen™ NuPAGE™ Sample Reducing Agent (Thermo Fisher Scientific), followed by incubation at 70°C for 10 min.

Protein samples were separated by SDS-PAGE using 4–12% Bis-Tris gels (Invitrogen) in Invitrogen™ NuPAGE™ MES SDS Running Buffer (Thermo Fisher Scientific). Proteins were subsequently transferred onto nitrocellulose membranes using the iBlot™ 2 Dry Blotting System (Thermo Fisher Scientific) according to the manufacturer’s instructions.

Primary and secondary antibodies were diluted in solutions from the iBind™ Solution Kit (Invitrogen). The following primary antibodies were used: rabbit anti-CASP3 (mouse and human; #9662, Cell Signaling Technology,), rabbit anti-CASP7 (human; #9492, Cell Signaling Technology), rabbit anti-CASP8 (mouse and human; #4790, Cell Signaling Technology), rabbit anti-FADD (mouse; #A307-154A, Antibodies.com), rabbit anti-FADD (human; #2782, Cell Signaling Technology), rabbit anti-Fas (human; #4233, Cell Signaling Technology), rabbit anti-Fas (mouse; #96825, Cell Signaling Technology), rabbit anti-GSDME (mouse; #ab215191, Abcam,), rabbit anti-MLKL (mouse; #37705, Cell Signaling Technology), rabbit anti-BID (mouse; #2003, Cell Signaling Technology), rabbit anti-BID (human; #2002, Cell Signaling Technology), rabbit anti-SEC62 (mouse; #ab140644, Abcam), and mouse anti-β-actin (mouse and human; #sc-47778, Santa Cruz Biotechnology). The secondary antibodies used were horseradish peroxidase (HRP)-conjugated anti-mouse IgG (#115-001-003) and anti-rabbit IgG (#111-001-003) antibodies (Jackson ImmunoResearch). Membranes were mounted onto iBind™ Cards wetted with iBind™ Solution and placed into an iBind™ Western Device (Invitrogen). Diluted primary and secondary antibodies were added according to the manufacturer’s protocol and incubated for a minimum of 2.5 h at room temperature (RT). Following antibody incubation, membranes were washed with deionized water for 5 min and incubated with SuperSignal™ West Pico PLUS Chemiluminescent Substrate (Thermo Fisher Scientific) for 2 min at RT. Chemiluminescent signals were detected using an iBright™ 1500 Imaging System (Invitrogen).

### Flow cytometry

WT and *Sec62^−/−^* P815 cells were washed with phosphate-buffered saline (PBS) and stained with Fixable Viability Dye Alexa Fluor™ 700 (Thermo Fisher Scientific) for 15 min at room temperature (RT). Following viability staining, cells were washed with PBS supplemented with 2% fetal bovine serum (FBS) and incubated with an APC-conjugated anti-Fas antibody (mouse; #17-0951-82, Thermo Fisher Scientific) for 30 min at RT. Cells were subsequently washed and fixed using Buffer A from the Human FoxP3 Buffer Set (BD Biosciences) for 10 min at RT. Following fixation, cells were resuspended in PBS containing 2% FBS and analyzed using a BD LSRFortessa™ flow cytometer (BD Biosciences). Data were analyzed using FlowJo v10.8.1 (BD).

### CRISPR–Cas9–mediated knockout generation

For the generation of knockout P815 cell lines, guide RNAs (gRNAs) targeting mouse genes were cloned into lentiCRISPR v2-derived lentiviral vectors carrying different antibiotic resistance markers: hygromycin (#98291, Addgene) for generation of *Casp3*⁻/⁻ cells, neomycin (#98292, Addgene) for generation of *Mlkl*⁻/⁻ cells, blasticidin (#98293, Addgene) for generation of *Bid*⁻/⁻ cells, and puromycin (#52961, Addgene) for generation of all other knockout cell lines. gRNAs were cloned according to the Zhang laboratory lentiCRISPR v2 cloning protocol ^56,57^.

Lentiviral particles were produced in Lenti-X™ 293T cells seeded at 5 × 10^5^ cells per well in 12-well plates and cultured in DMEM supplemented with 10% FBS and 20 mM HEPES. Cells were co-transfected with the lentiviral transfer vector (1.33 µg/well), the packaging plasmid psPAX2 (1.0 µg/well), and the envelope plasmid pCMV-VSV-G (0.44 µg/well) using Lipofectamine™ 2000 (Invitrogen; 5 µL/well) in 200 µL Opti-MEM™ (Gibco). The culture medium was replaced 24 h after transfection. Viral supernatants were harvested 72 h post-transfection, clarified by centrifugation, and 300 µL of viral supernatant was used to transduce P815 cells (5 × 10⁴ cells/well in 12-well plates). Following transduction, cells were selected with the appropriate antibiotic to generate stable gRNA-expressing cell populations. Antibiotic-resistant cells were seeded into 96-well plates at a density of 0.75 cells/well to obtain single-cell-derived clones. Individual colonies were expanded and screened for loss of target protein expression by Western blot analysis.

For generation of knockout HeLa^EGFP^ cell lines, gRNAs targeting human genes were cloned into the pSpCas9(BB)-2A-Puro (PX459) V2.0 vector (#62988, Addgene) according to the Zhang laboratory cloning and target sequencing protocol ^58^. HeLa^EGFP^ cells were seeded at 1 × 10⁵ cells/well in 12-well plates and transfected with 1.6 µg of gRNA-containing plasmid using Lipofectamine™ 2000 (5 µL/well) in 200 µL Opti-MEM™. Twenty-four hours after transfection, cells were subjected to puromycin selection for 72 h. Surviving cells were expanded in T25 flasks for 48 h before being seeded into 96-well plates at a density of 0.75 cells/well. Single-cell-derived colonies were isolated, expanded, and screened for loss of target protein expression by Western blot analysis.

DNA sequences used for cloning of gRNAs in this study are summarized in **Supplementary Table III**.

### Genome-wide CRISPR screen

The mouse genome-wide Brie CRISPR knockout pooled library (lentiGuide-Puro, #73633, Addgene) was amplified using Endura™ ElectroCompetent Cells (Biosearch Technologies; #60242-2) according to the manufacturer’s instructions. Colony numbers were determined from serial dilutions to ensure ≥100-fold representation of the library diversity (4 sgRNAs per gene). Colonies were harvested by scraping in LB medium, and plasmid DNA was extracted using an endotoxin-free maxiprep kit (NucleoBond Xtra Maxi Plus EF) and stored at −20°C.

For the genome-wide screen, 7.5 × 10⁷ *Casp3⁻/⁻* (hygromycin-resistant) or *Casp3⁻/⁻Mlkl⁻/⁻Bid⁻/⁻* (hygromycin-, neomycin-, and blasticidin-resistant) P815 cells were transduced with the Brie lentiviral library at a multiplicity of infection (MOI) of 0.3 to ensure single integration events per cell. 48 hours post-transduction, cells were selected with puromycin (0.6 µg/mL). After 48 h of selection, surviving cells were pooled, cultured in puromycin-free medium, and subjected to CTL selection.

For CTL selection, 8 × 10⁶ target cells were co-cultured with activated human CD8⁺ T cells at equal effector-to-target ratios in the presence of 5 µg/mL anti-CD3 antibody (#317347, BioLegend). Four biological replicates were used for *Casp3⁻/⁻* cells and three replicates for *Casp3⁻/⁻Mlkl⁻/⁻Bid⁻/⁻* cells. After 24 h (*Casp3⁻/⁻*) or 48 h (*Casp3⁻/⁻Mlkl⁻/⁻Bid⁻/⁻*) of co-culture, cells were collected and treated with puromycin to eliminate residual CD8⁺ T cells. Surviving P815 cells were expanded, and CTL selection was repeated three times for *Casp3⁻/⁻* cells and twice for *Casp3⁻/⁻Mlkl⁻/⁻Bid⁻/⁻* cells. Following selection, surviving cells were expanded, and genomic DNA was extracted from 2 × 10⁷ cells per replicate using the NucleoSpin™ Blood L Midi kit (Macherey-Nagel; #740954.20). sgRNA sequences were amplified and prepared for next-generation sequencing (NGS) using a two-step PCR strategy. In the first PCR (30–45 cycles), sgRNA-containing regions were amplified from genomic DNA using eight staggered forward primers (PCR1_Fwd_1–8) together with a vector-specific reverse primer, using Q5® High-Fidelity DNA Polymerase (New England Biolabs; M0491S). PCR products were pooled and purified using the NucleoSpin™ Gel and PCR Clean-up Midi kit (Macherey-Nagel; #740986.20). In the second PCR (8 cycles), Illumina P5 and P7 adapters and sample-specific indices were added. Final libraries were purified using SpeedBeads (GE65152105050250, Cytiva), quantified using a Qubit fluorometer (Thermo Fisher Scientific), and quality-checked using TapeStation system with D1000 ScreenTape (5067-5582, Agilent). Libraries were pooled equimolarly and sequenced on a NovaSeq X platform (Illumina), targeting a depth of 20–30 million reads per sample.

Screening data were analyzed using MAGeCK-VISPR ^59^ to identify significantly enriched or depleted sgRNAs/genes. In brief, sgRNA counting was first performed using the *mageck count* function on FASTQ files. CTL–treated and non-treated samples were then compared using the *mageck test* function, and normalization was performed using control sgRNAs.

The primers used for the first-round PCR are listed below. Blue indicates the sequences used for amplification of the gRNA within the lentiGuide-Puro vector backbone, and red indicates stagger sequences introduced to increase sequence diversity.

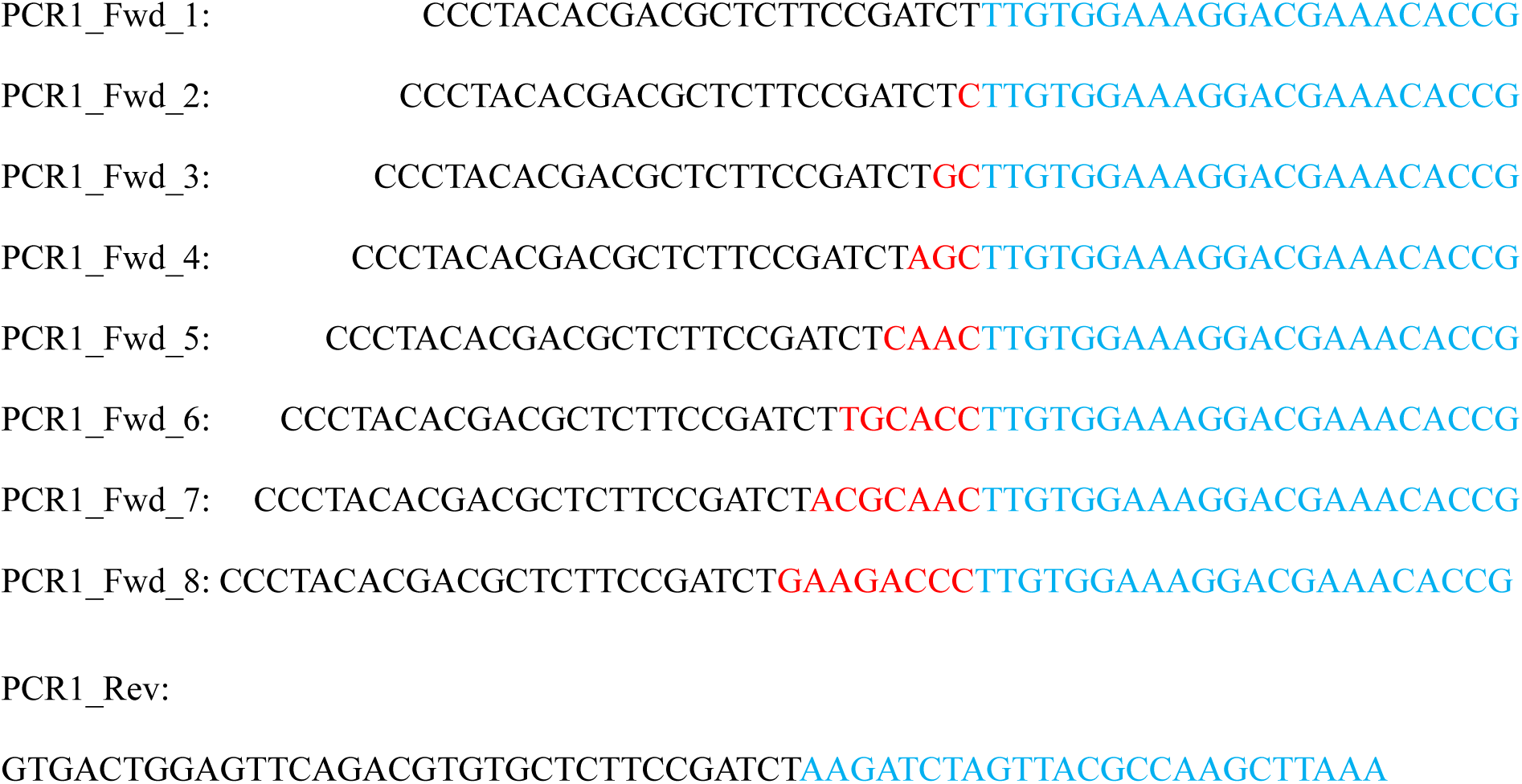

Primers used for the second PCR were as follows, where N₈ represents sample-specific index sequences:

Forward: AATGATACGGCGACCACCGAGATCTACACNNNNNNNNACACTCTTTCCCTACACGACGCTCTTC CGATC*T

Reverse: CAAGCAGAAGACGGCATACGAGATNNNNNNNNGTGACTGGAGTTCAGACGTGTGCTCTTCCG ATC*T

### CTL–αCD3–P815 killing assay

P815^ZsGreen1^ cells were prepared at a concentration of 10⁵ cells/mL in P815 culture medium. Anti-CD3 antibody (#317347, BioLegend) was added to a final concentration of 3.6 µg/mL, and cells were seeded into 96-well plates (100 µL per well). Each condition (with or without CTLs) was performed in three technical replicates. Cells were incubated for 1 h at 37 °C in 5% CO₂ prior to the addition of activated CD8⁺ CTLs (100 µL per well; 10⁵ cells/mL in CD8⁺ culture medium, with beads removed using a magnetic stand), corresponding to an effector-to-target ratio of 1:1. Plates were then placed in an Incucyte® S3 or S5 live-cell imaging system (Sartorius) and imaged every 2 h for 72–96 h.

### NK-92–HeLa killing assay

HeLa^EGFP^ cells were seeded at 2 × 10⁴ cells/mL in HeLa culture medium and allowed to adhere for 24 h. Each condition (with or without NK-92 cells) was performed in three technical replicates. NK-92 cells were prepared in NK-92 culture medium and added to HeLa cultures at an effector-to-target ratio of 1:1 or 3:1, assuming approximately twofold expansion of HeLa cells over 24 h. Plates were then incubated in an Incucyte® S3 or S5 live-cell imaging system (Sartorius) for 46–50 h, with images acquired every 2 h.

### TCGA database analysis

The number and frequency of simple somatic mutations (SSMs), excluding copy number variations, for individual genes were manually extracted from The Cancer Genome Atlas (TCGA) by querying genes of interest through the Genomic Data Commons Data Portal (https://portal.gdc.cancer.gov/). A pan-cancer dataset was generated by aggregating data across all TCGA projects without stratification by tumor type. Variant annotations, including Ensembl Variant Effect Predictor (VEP) ^49^ classifications, were manually extracted from TCGA and used for data visualization. Following definitions are used for the impact ratings:

- HIGH (HI): The variant is assumed to have high (disruptive) impact in the protein, probably causing protein truncation, loss of function or triggering nonsense mediated decay: *transcript_ablation, splice_acceptor_variant, splice_donor_variant, stop_gained, frameshift variant, stop_lost, start_lost, transcript_amplification*.
- MODERATE (MO): A non-disruptive variant that might change protein effectiveness: *inframe_insertion, inframe_deletion, missense_variant, protein_altering_variant, regulatory_region_ablation*.
- LOW (LO): Assumed to be mostly harmless or unlikely to change protein behavior: *splice_region_variant, incomplete_terminal_codon_variant, stop_retained_variant, synonymous_variant*.
- MODIFIER (MR): Usually non-coding variants or variants affecting non-coding genes, where predictions are difficult or there is no evidence of impact: *coding_sequence_variant, mature_miRNA_variant, 5_prime_UTR_variant, 3_prime_UTR_variant, non_coding_transcript_exon_variant, intron_variant, NMD_transcript_variant, non_coding_transcript_variant, upstream_gene_variant, downstream_gene_variant, TFBS_ablation, TFBS_amplification, TF_binding_site_variant, regulatory_region_amplification, feature_elongation, regulatory_region_variant, feature_truncation, intergenic_variant*.

### Statistical analysis

Statistical analyses were performed using GraphPad Prism version 10.6.1. A two-tailed unpaired *t*-test was used for comparisons between two groups, while one-way ANOVA was used for comparisons among more than two groups. Differences were considered statistically significant as indicated in the figure legends.

## Supporting information

Supplementary Table III

## Data Availability statement

The CRISPR screen datasets have been deposited in the NCBI Gene Expression Omnibus (GEO) under accession numbers GSE336303 and GSE336305. Additional datasets generated and/or analyzed during the current study are available from the corresponding authors upon reasonable request.

## Author Contributions

E.S.: Methodology, Data curation, Visualization, Writing – original draft, review, and editing. S.W.: Conceptualization, Methodology, Data curation, Visualization, Writing – review and editing. Q.W.: Methodology, Data curation, Writing – review and editing. N.E.B.S.: Methodology, Writing – review and editing. K.T.: Supervision, Project administration, Resources, Conceptualization, Writing – review and editing. Y.L.: Supervision, Project administration, Resources, Conceptualization, Methodology, Data curation, Visualization, Writing – original draft, review, and editing.

## Acknowledgements

We thank Mentowa Fürst Bright, Nora Rojahn Bråthen, Jenny Karoline Jebsen and Siri Sæterstad for their expert technical assistance. We are grateful to the Norwegian Sequencing Centre for assistance with CRISPR screen sequencing. The illustration in Figure 1A was created using BioRender (https://www.biorender.com).

This work was supported by the Research Council of Norway (grant 344942 to Y.L. and grant 328827 to KT), the Astri and Birger Torsteds foundation (to Y.L.), the Norwegian Cancer Society (grant 298910 to K.T.) and the Regional Health Authority for South-Eastern Norway (grant 2025081 to KT). Y.L. also received funding from the European Union’s Horizon 2020 Research and Innovation Programme through the Marie Skłodowska-Curie Actions (Scientia Fellowship, Grant Agreement No. 801133).

